# Discovery of dual inhibitors of KPC-3 and KPC-15 of Klebsiella pneumoniae – an in-silico molecular docking and dynamics study

**DOI:** 10.1101/2019.12.23.886671

**Authors:** Sony Sharma, Praveen Kumar-M, Santa Singh, Ajay Sohal, Rohitash Yadav, Nitin Gupta

## Abstract

**Background:** The development of carbapenem resistance against Klebsiella pneumoniae is a situation of grave concern and requires urgent attention. Among the KPC produced by K.pneumoniae, KPC-3, and KPC-15, play a significant role in the development of resistance to carbapenem.

**Materials and methods:** The binding sites of KPC-3 and KPC-15 were predicted by the COACH server. Drug-like ligands from ZINC were then screened by ligand-based drug screening (LBVS) by keeping Relebactam as a template. The top 50,000 selected ligands were then screened by structure-based virtual screening using idock. For keeping an account of the dual inhibitors’ stability in complex with KPC-3 and KPC-15, MDS were carried out for each complex.

**Results:** Based on consensus weighted ranks, the top 3 ligands with the dual inhibitory property are ZINC76060350 (consensus weighted rank - 1.5), ZINC05528590 (2), ZINC72290395 (3.5). All the top 3 dual inhibitors have a reasonable probability of passing through the blood-brain barrier. The RDKit and Morgan fingerprint scores between Relebactam and the top three ligands were 0.24, 0.22, 0.23, and 0.26, 0.19, 0.25, respectively (showing only 20% similarity). The MD simulation result revealed good binding stability of ligand ZINC05528590 with both KPC-3 and KPC-15, whereas ligand ZINC76060350 showed good binding stability to KPC-3.

**Conclusion:** The ligand ZINC05528590 could be taken forward to develop a new drug against a multi-resistant- Klebsiella pneumoniae infection. At the same time, ZINC76060350 can be considered to develop a new drug against KPC-15 resistant Klebsiella pneumoniae.

## Introduction

Carbapenem antibiotics, such as imipenem and meropenem, have been one of the best available antibiotics, which are used as the first line of therapy in case of severe infection caused by extended-spectrum beta (b)-lactamases (ESBL) producing Enterobacteriaceae [1–3]. The activity of carbapenem could be augmented by the addition of beta-lactamase inhibitors. An example of this is imipenem, which showed a significant improvement in activity against various species of Enterobacteriaceae following the addition of Relebactam [as assessed by a decrease in the minimum inhibitory concentration (MIC)] [4]. Carbapenem-resistant Enterobacteriaceae is a grave concern against this potent drug, and it is spreading rapidly all around the globe with an increase in morbidity and mortality associated with healthcare-associated infection (HAI). The mechanism involved in the development of resistance to carbapenem is multi-fold, ranging from (i) modifications in outer membrane permeability, (ii) the up-regulation of the efflux system corresponding to the hyperproduction of AmpC β lactamases (cephalosporinases) or ESBL’s, and (iii) the production of an enzyme which degrades the carbapenem (carbapenemases) [5,6]. Among the Carbapenem-resistant Enterobacteriaceae, the most common bacteria, is K. pneumoniae [7].

Klebsiella pneumonia, a gram-negative nosocomial bacterium, is responsible for both community-acquired infection and HAI [8]. Carbapenem acts by binding to the penicillin-binding proteins (PBP) of the bacteria, thereby resulting in the bacterial cell wall’s lysis. The binding will instigate the bacterial cell to produce carbapenemase, which binds with the carbapenems to hydrolyze the antibiotics containing beta-lactams [9]. Carbapenemases (KPC-2 & it’s variant KPC-3 and KPC – 15) production by K. pneumoniae is mainly responsible for the resistance against carbapenem [10–13]. Based on the molecular classification, the carbapenemases are divided into classes A to D. In classes A and C; the active sites contain serine (non-metallo), referred to as non-metallo β-lactamases. In contrast, class B contains zinc (metal) and is referred to as metallo β-lactamases [8]. Class A β-lactamases comprise carbapenemases such as NMC, IMI, SME, GES and, KPC (variants from KPC-2 to KPC-17) families. In contrast, class D consists of β-lactamases, which are of oxacillinase variety. Class B metallo β-lactamases contain IMP, GIM, VIM, SPM, and SIM [14]. KPC-3 and KPC-15 are two of the most frequently found carbapenemases and hydrolyze a wide variety of β-lactam antibiotics [8,15]. Various reports indicate that KPC-15 also shows resistance to almost 18 conventional antimicrobial agents, including carbapenem antibiotics [12].

With the current advancement in bioinformatics, Computer-Aided Drug Discovery (CADD) offers multiple-tools to make drug discovery affordable at a reduced cost. In addition to the efficacy of compounds, ADMET characteristics also affect drugs development [16,17]. Hence, in this study, we aspired to identify a dual inhibitor of KPC-3 and KPC-15 by employing Ligand Based Virtual Screening (LBVS), Structure-Based Virtual Screening (SBVS), the evaluation of ADMET properties of top compounds, and finally confirming our findings by Molecular Dynamics (MD).

## Material and Methodology

A workstation - HP-Z640 working on Ubuntu 18.04.2 LTS with a configuration of Intel Xeon(R) CPU E5-2640 v3@2.60 GHz × 16 (hyperthreaded cores), 64 GB RAM, 500 GB HDD and Quadro K620/PCLe/SSE2 graphics card was used for conducting all the studies. Open source in-silico offline tools and webservers (or) proprietary tools with appropriate academic licenses were used for conducting the experiments. Their reason for selection has been given in the respective section of methodology. Broadly the identification of active ligands against KPC-3 and KPC-15 can be divided into ten steps, namely: 1. Retrieval of ligands; 2. Ligand-based virtual screening; 3. The accession of target protein; 4. Preparation of receptor for docking; 5. Active site prediction; 6. Structure-based virtual screening; 7. Analysis of docking results; 8. Structure similarity search; 9. Determination of ADMET properties; 10. MD Simulation

### 1. Retrieval of the ligands

“Drug-like” molecules subset within the ZINC12 database (http://zinc.docking.org) were used for the retrieval of ligands. The downloaded library consisted of around 1 million ligands in 5 set in mol2 format (0.2 million ligands in each set). These being “Drug-like” molecules by default obey Lipinski’s Rule of Five as well as follow specific criteria of drug-likeness, namely high potency, ligand efficiency, lipophilic efficiency and bioavailability.

### 2. LBVS

LBVS uses the information of known active ligand for prediction. It does not require the knowledge of target protein structure for screening [18]. The ligand-based tool known as LiSiCA was used to screen all five sets of ligands retrieved from the previous step. LiSiCA uses a clique algorithm to find two- and three- dimensional similarities between pairs of ligands as supplied in mol2 atom types [19]. Relebactam being an already known inhibitor of KPC, was taken as the active reference ligand for both KPC-3 and KPC-15 structure [20]. The following input parameters were used while running LiSiCA (i) dimension (d) was 3, (ii)-m=0.5 for Rigorous screening, (iii)-c=10000 for a maximum number of output files. LiSiCA. The resulted output files contained 10000 ligands per set giving a pool of 50000 compounds. These resulted output files obtained from LiSiCA (mol2) were converted into pdbqt format using a python script supplied with MGLtools (prepare_ligand4.py). These top 50000 ligands were taken for SBVS using idock.

### 3. The accession of the target protein

RCBS protein data bank, a global archive of protein and nucleic acids structures in 3D the format was used for retrieving the protein target structure. As the 3D structure of KPC-3 and KPC-15 was not available in the RCBS protein data bank (at the time of screening - April 2019), we took the structure of KPC-2 carbapenemase from K. pneumoniae (PDB ID 2OV5) and decided to mutate the structure to arrive at KPC-3 and KPC-15. The structure obtained of KPC-2 was the X-ray crystallographic structure of 1.85 Å resolution with no ligands attached. Chain A of protein was separated using pymol [21]. This file was exported for the further mutation to obtain KPC-3 (H272Y) and KPC-15 (A119L and G146K) [12] using Swiss PDB Viewer [22] in PDB format.

### 4. Preparation of the receptor for docking

MGLTools inside AutoDock were used for preparing the receptor. The water molecules of the target protein were deleted for the computational convenience of bioenergies and for clearing the active binding pockets, which could potentially distort the pose search. The AutoDock uses the United Atom force field of AMBER, which only uses polar hydrogens atoms. Therefore, we added only polar hydrogen atoms and merged the non-polar hydrogen atoms to get the correct ionization state and tautomer [23,24]. Also, it helps in reducing the number of atoms to be modeled explicitly during docking. Finally, the Kollman United Atom charges were assigned to the protein structure. The modified protein was then converted into a pdbqt structure.

### 5. Active site prediction

The COACH server was used to estimate the active site of the protein (Yang et al., 2013a). The PDB file of KPC-3 and KPC-15 was uploaded as a query in the COACH server, and it took approximately 10 hours to predict the active site coordinates. The summary of COACH was computed based on the results of TM-SITE, S-SITE, COFACTOR, FINDSITE, ConCavity. The final summary consisted of residue-specific binding probability, template clustering results, and predicted bound ligands. It also provides consensus binding residues between the uploaded protein and the PDB hit along with their respective C-score values and cluster size.

### 6. SBVS

In SBVS, a library of ligands was docked into an identified docking site of the protein. We used Autodock Vina for detecting the biophysical interaction between ligand and protein [25]. idock is an improvement over the AutoDock Vina and is preferred for 3 important reasons: it enables the process of multi-threading in a multi-core computer; permits the storage of details about grid maps and the receptor in the working memory - thereby avoiding repetitive detection for each ligand, enables the detection and avoidance of ligands having inactive torsions. All the above factors significantly improve and reduce the time required for ligand-protein docking [26,27]. Behind idock is an algorithm based on regression and optimization theorems giving an estimate of the biophysical interaction in terms of binding energy (or) idock score. The lower the binding energy, the more is the interaction between the ligand and protein, and this helps in ranking and identifying the active ligand. Also, confirmation of the ligand at the docked site and the plausibility of the given structure of the ligand are taken into consideration during docking. A shell script was coded for repeating the process of docking, and the final outputted results were ranked based on binding energy for further analysis. The dual inhibitors of both KPC-3 and KPC-15 were identified based on consensus weighted rank. Consensus weighted rank was calculated by dividing the mean of the ranks of the compound observed in docking against each protein by the predicted number of proteins inhibited by the molecules. Based on the initial results of docking of Relebactam KPC-3 and KPC-15, we took a cut-off of −6 kcal/mol binding energy (both KPC-3 and KPC-15) for identifying a molecule as an inhibitor of a protein (any ligand showing binding energy of less than −6 kcal/mol was taken as inhibitor). The lower the consensus weighted rank more potent is the dual inhibitor.

### 7. Analysis of the docking results

PymoL was used for the preparation of complex PDB file of protein and identified ligands. It was also used for analyzing the docking position using the same. Ligplot+ was used for studying the interaction between the ligand and protein and for identifying the bond length [28].

### 8. Structure similarity search

Structure similarity between relebactam and top 3 identified ligands (dual inhibitors) were made using the python package RDKit [29]. The canonical SMILES were used as an input query and the depicted 2D structures were converted into 3-D structures using distance geometry for structure comparison. RDKit performs structure comparison using two prominent fingerprints (i) RDKit fingerprint, for substructure fingerprinting where the atom types are set by atomic number and aromaticity; bond types based on atom types and bond types (ii) Morgan fingerprint is a reimplementation of the extended connectivity fingerprint (ECFP), which takes into account the neighborhood of each atom for similarity fingerprint.

### 9. Determination of ADMET properties

We employed admetSAR to further study the ADMET properties of the identified lead ligands. The Forty endpoints of ADMET properties were identified using the admetSAR tool, which takes the input in the form of canonical SMILES. [17].

### 10. MD Simulation for protein-ligand complex

The protein-ligand complex was further evaluated based on their structural stability by Molecular Dynamics Simulation (MDS) using GROMACS 2020. [30]. The stability of top two dual inhibitors was evaluated individually with KPC-3 and KPC-15 using MD Simulation. First, we generated a topology file using the SwissParam webserver. [31]. CHARMM36 (March 2019) was used as a force field. The solvation for the protein-ligand complex was made using TIP3P explicit water molecule in a triclinic shaped box. Sodium (Na+) and Chlorine (Cl) ions were added to neutralize the system’s net charge substituted by solvent molecules in the system. Steepest Descent algorithm was used for the energy minimization of the system with a tolerance of 1000 kJ/mol/nm. Periodic Boundaries and Van der Walls cut-off of 12 A were assigned in all directions. NVT and NPT ensemble MD simulation was run for 100 ps in the periodic boundary condition. Modified Berendsen thermostat & barostat were used to keep the temperature constant at 300K and pressure at 1 bar. Long-Range Electrostatic interaction was calculated by implying the Particle Mesh Ewald method. LINCS Algorithm was used for constraining the bond lengths. The temperature and pressure were kept constant during MD simulation for 10 ns and 20 ns for the ligands. The final run was used to analyze root mean square deviation (RMSD) of protein (Backbone, for least-square fitting) and ligand, root mean square fluctuation (RMSF), and the number of hydrogen bonds.

## Results and Discussion

### 1. Identification of ligand binding sites

The COACH and COFACTOR were used to identify the active sites of KPC-3 and KPC-15 of Klebsiella pneumoniae via the COACH server. The results obtained from the COACH showed a close relation with 2jbfA (chain A of PDB ID 2JBF) with a C-score (confidence score of the predicted result) value of 1.00 for KPC-3, while for KPC-15, the close relation was observed with 4xuzA (chain A of PDB ID 4XUZ) with a C-score value of 0.87. It ranges between 0-1 (inclusive of the endpoints), where the higher C-score value indicates a more reliable prediction. The hydrolysis of the substrate increases with the increase in the interaction between the substrate and the beta-lactamase. For both KPC-3 and KPC-15, the COFACTOR represented a close match with 2ov5A (chain A of PDB ID 2OV5) with a TM score (template modelling score - estimates the protein structure similarity) equals to 1.00 and root mean square deviation (RMSD) equals to 0.00 Å. With a smaller value of RMSD, a higher resolution of the protein structure is observed [32]. Like C-score, TM also ranges between 0-1 (inclusive of the endpoints), where a higher value indicates a similarity between structures. The predicted putative ligand binding site residues in KPC-3 are- Cys39, Ser40, Lys43, Trp75, Ser100, Asn102, Leu137, Asn140, Thr186, Lys204, Thr205, Gly206, Thr207 while in KPC-15 are- Cys39, Ser40, Lys43, Pro74, Trp75, Ser100, Asn102, Leu137, Asn140, Lys204, Thr205, Gly206, Thr207, Cys208, Val210. The final COACH results, (i.e.,) the predicted active site coordinates for both KPC-3 and KPC-15 were- x = 55.891, y = −21.594, z = −4.513.

### 2. Virtual screening and analysis

The top 3 potential inhibitor of KPC-3 as predicted by idock are ZINC05528590 (ligand 1a, −6.41 kcal/mol), ZINC76060350 (ligand 2a, −6.37 kcal/mol), ZINC72290395 (ligand 3a, −6.33 kcal/mol) (Table 1). The best ranked compound, ZINC05528590, IUPAC name- (5S)-5-hydroxy-5-methyl-1-oxa-3,4- diazaspiro[5.5]undecan-2-one, has a molecular weight of 200.23g/mol with H-bond donor of 3 and H-bond acceptor of 4. The top 3 potential inhibitor of KPC-15 as predicted by idock are ZINC76060350 (ligand 1b, −6.39 kcal/mol), ZINC22689973 (ligand 2b, −6.38 kcal/mol), ZINC05528590 (ligand 3b, −6.34 kcal/mol) (Table 2). ZINC76060350 with an IUPAC name of (3R)-1-[(1R)-cyclohex-2-en-1-yl]-3-(5-methyl-1H-pyrazol-3-yl) piperidine having a molecular weight of 245.36g/mol, showed best docking score and having 1 & 2 H-bond donors and acceptors, respectively. Based on consensus weighted ranks, the top 3 ligands with the dual inhibitory property are-ZINC76060350 (ligand 1c, consensus weighted rank - 1.5), ZINC05528590 (ligand 2c, consensus weighted rank - 2), ZINC72290395 (ligand 3c, consensus weighted rank - 3.5) (Table 3). The binding of the ligand 1c with the active site of KPC-3 and KPC-15 along with the hydrogen bond interaction with bond length are shown in figure 1a and figure 1b, respectively. The LigPlot+ images of the top 3 dual inhibitors of KPC3 and KPC15 are denoted in figure 2. All the selected compounds showed an overall high binding affinity towards the target protein, i.e. (the minimum binding the energy of the 10th rank ligand against KPC-3 was −6.22 kcal/mol, and KPC-15 was - 6.17 kcal/mol).

**Table 1:**
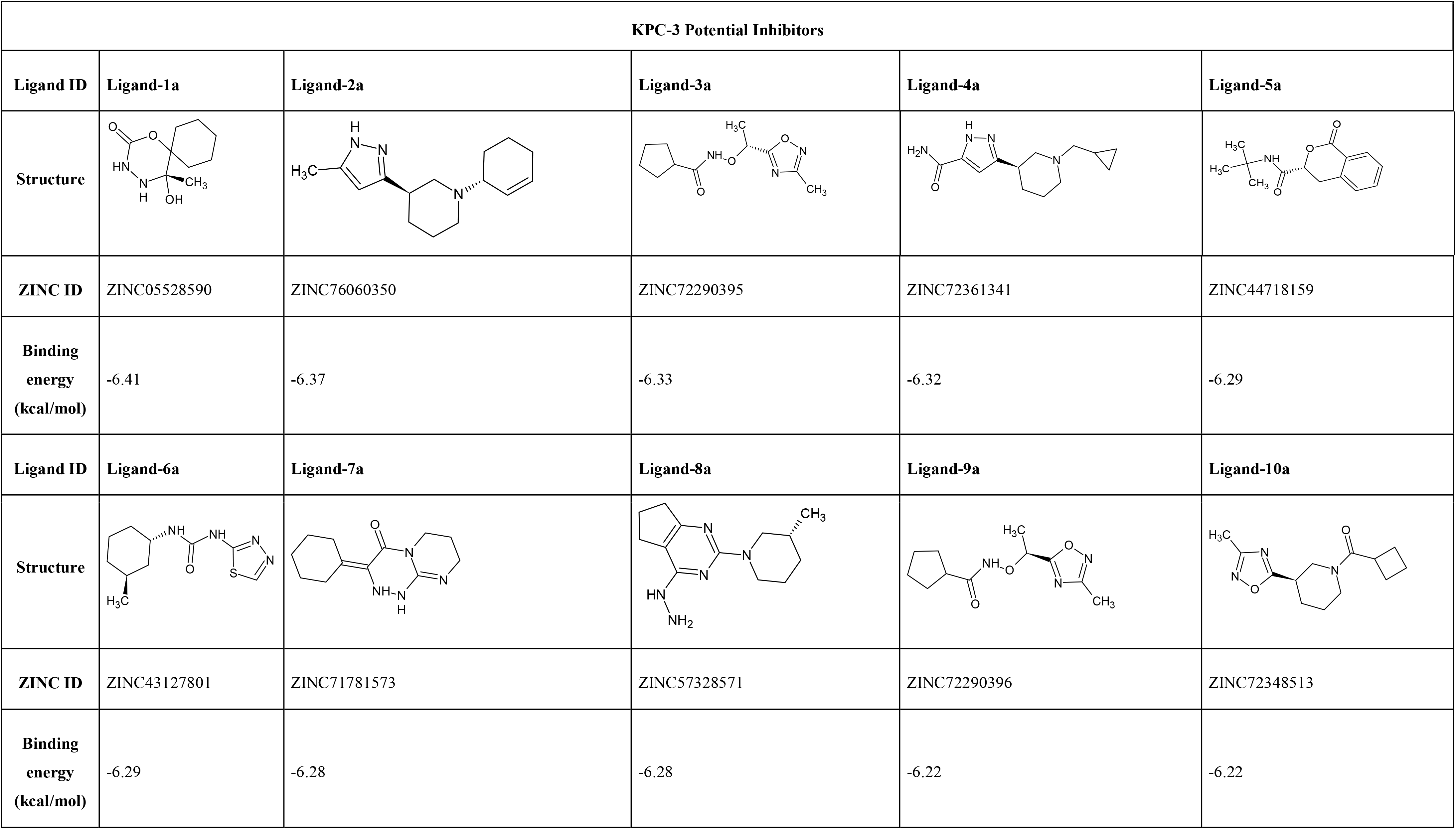
The top 10 potential inhibitors against KPC-3 are reported here according to the lowest binding energy. The Ligand-1a is the best compound seen with the value of −6.41, ID: ZINC05528590, IUPAC name- (5S)-5-hydroxy-5-methyl-1-oxa-3,4-diazaspiro[5.5]undecan-2-one

| KPC-3 Potential Inhibitors |  |  |  |  |  |
| --- | --- | --- | --- | --- | --- |
| Ligand ID | Ligand-1a | Ligand-2a | Ligand-3a | Ligand-4a | Ligand-5a |
| Structure |  |  |  |  |  |
| ZINC ID | ZINC05528590 | ZINC76060350 | ZINC72290395 | ZINC72361341 | ZINC44718159 |
| Binding energy (kcal/mol) | -6.41 | -6.37 | -6.33 | -6.32 | -6.29 |
| Ligand ID | Ligand-6a | Ligand-7a | Ligand-8a | Ligand-9a | Ligand-10a |
| Structure |  |  |  |  |  |
| ZINC ID | ZINC43127801 | ZINC71781573 | ZINC57328571 | ZINC72290396 | ZINC72348513 |
| Binding energy (kcal/mol) | -6.29 | -6.28 | -6.28 | -6.22 | -6.22 |

**Table 2:**
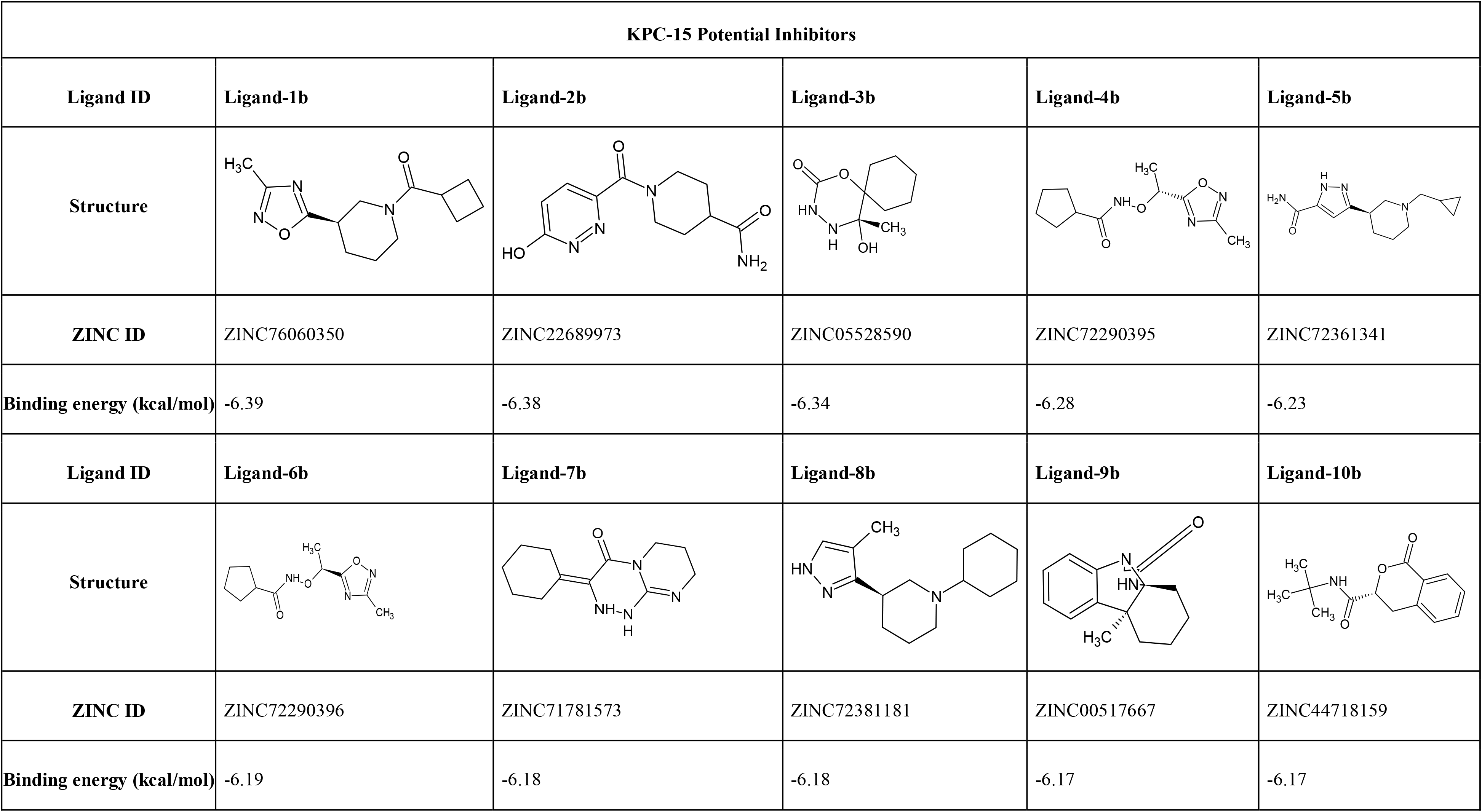
The top 10 potential inhibitors against KPC-15 are reported here according to the lowest binding energy. The Ligand-1b is the best compound seen with the value of −6.39, ID: ZINC76060350, IUPAC name- (3R)-1-[(1R)-cyclohex-2-en-1-yl]-3-(5-methyl-1H-pyrazol-3-yl)piperidine.

| KPC-15 Potential Inhibitors |  |  |  |  |  |
| --- | --- | --- | --- | --- | --- |
| Ligand ID | Ligand-1b | Ligand-2b | Ligand-3b | Ligand-4b | Ligand-5b |
| Structure |  |  |  |  |  |
| ZINC ID | ZINC76060350 | ZINC22689973 | ZINC05528590 | ZINC72290395 | ZINC72361341 |
| Binding energy (kcal/mol) | -6.39 | -6.38 | -6.34 | -6.28 | -6.23 |
| Ligand ID | Ligand-6b | Ligand-7b | Ligand-8b | Ligand-9b | Ligand-10b |
| Structure |  |  |  |  |  |
| ZINC ID | ZINC72290396 | ZINC71781573 | ZINC72381181 | ZINC00517667 | ZINC44718159 |
| Binding energy (kcal/mol) | -6.19 | -6.18 | -6.18 | -6.17 | -6.17 |

**Table 3:**
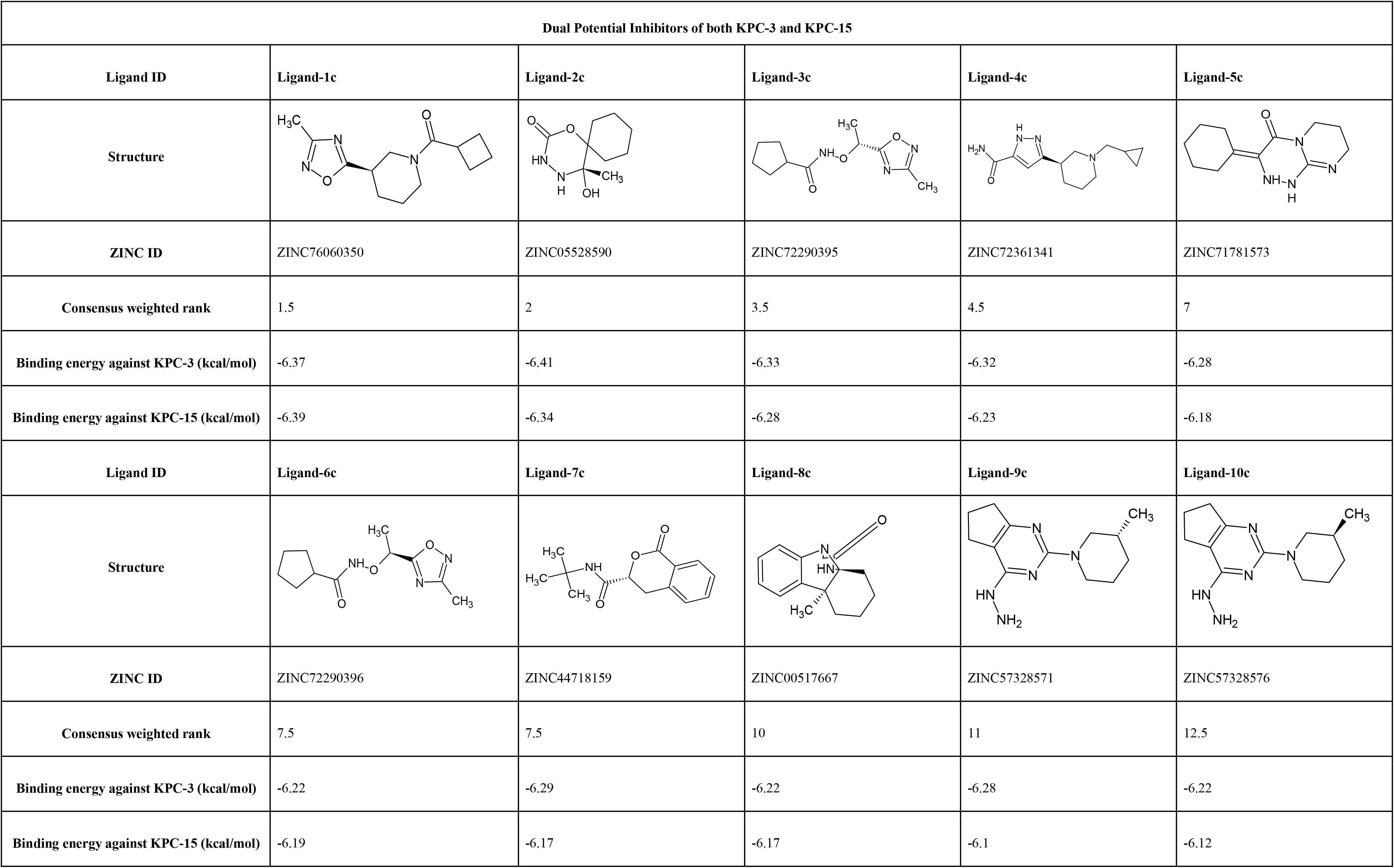
The top 10 potential inhibitors against KPC-3 and KPC-15 are reported here according to the lowest binding energy. The Ligand-1c is the best compound seen with consensus weighted rank of 1.5, ID: ZINC76060350, IUPAC name- (3R)-1-[(1R)-cyclohex-2-en-1-yl]-3-(5-methyl-1H-pyrazol-3-yl)piperidine.

| Dual Potential Inhibitors of both KPC-3 and KPC-15 |  |  |  |  |  |
| --- | --- | --- | --- | --- | --- |
| Ligand ID | Ligand-1c | Ligand-2c | Ligand-3c | Ligand-4c | Ligand-5c |
| Structure |  |  |  |  |  |
| ZINC ID | ZINC76060350 | ZINC05528590 | ZINC72290395 | ZINC72361341 | ZINC71781573 |
| Consensus weighted rank | 1.5 | 2 | 3.5 | 4.5 | 7 |
| Binding energy against KPC-3 (kcal/mol) | -6.37 | -6.41 | -6.33 | -6.32 | -6.28 |
| Binding energy against KPC-15 (kcal/mol) | -6.39 | -6.34 | -6.28 | -6.23 | -6.18 |
| Ligand ID | Ligand-6c | Ligand-7c | Ligand-8c | Ligand-9c | Ligand-10c |
| Structure |  |  |  |  |  |
| ZINC ID | ZINC72290396 | ZINC44718159 | ZINC00517667 | ZINC57328571 | ZINC57328576 |
| Consensus weighted rank | 7.5 | 7.5 | 10 | 11 | 12.5 |
| Binding energy against KPC-3 (kcal/mol) | -6.22 | -6.29 | -6.22 | -6.28 | -6.22 |
| Binding energy against KPC-15 (kcal/mol) | -6.19 | -6.17 | -6.17 | -6.1 | -6.12 |

**Figure 1:**
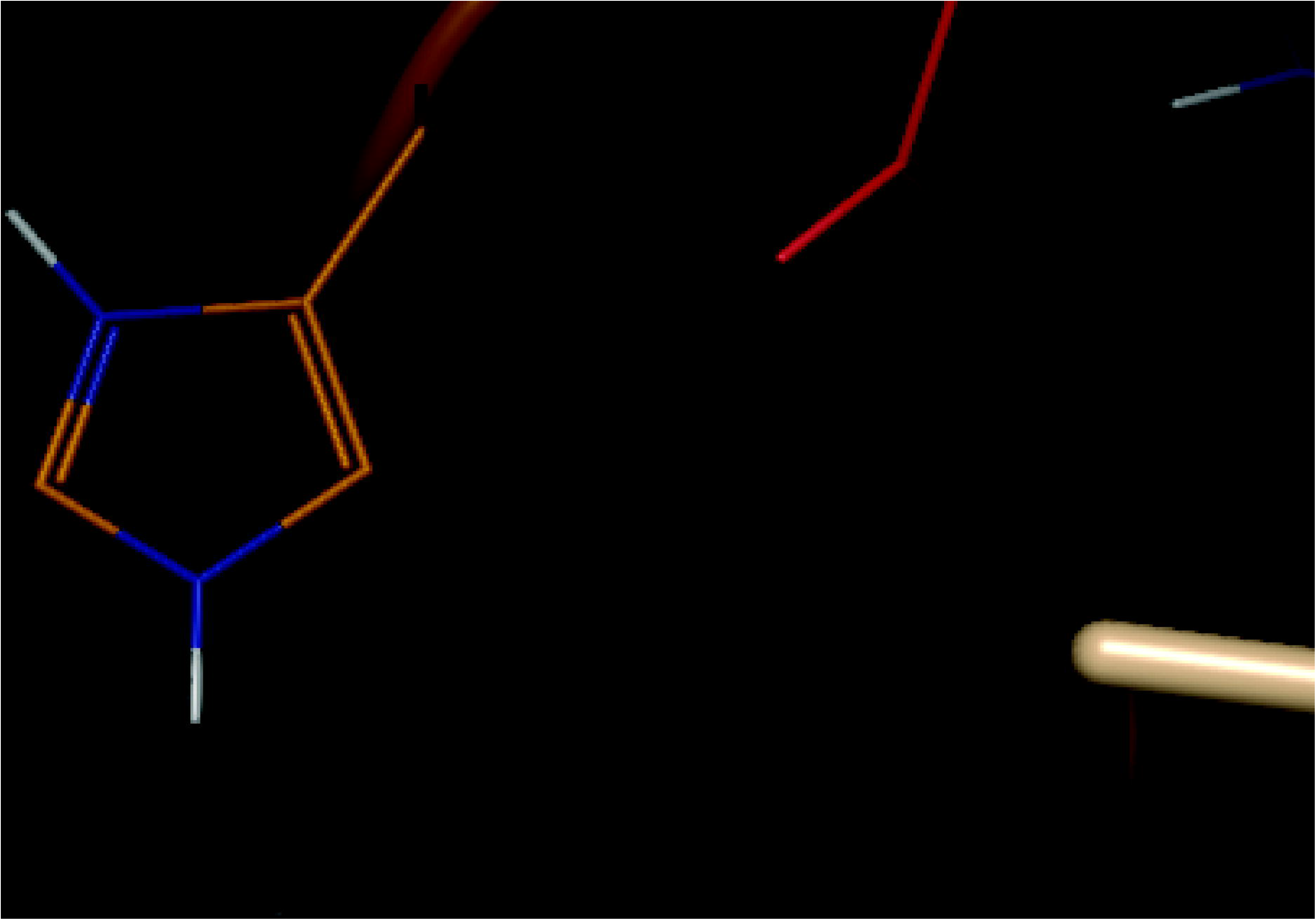
A) Interaction of KPC-3 to ZINC05528590 with a binding energy of −6.41 kcal/mol. B) Interaction of KPC-15 to ZINC76060350 with a binding energy of −6.39 kcal/mol. The top part of the figure shows the surface representation of the protein and ligand in ball & stick appearance whereas the bottom part of the figure is the zoomed in binding site with helical representation of the protein and ball & stick appearance of the ligand. The dotted yellow line represents the hydrogen bond interaction between the ligand and the protein. In case of KPC-3 there are 2 hydrogen bonds (2.3 Å and 3.2 Å), while incase of KPC-15 there is 1 hydrogen bond (2.3 Å) interaction.

**Figure 2:**
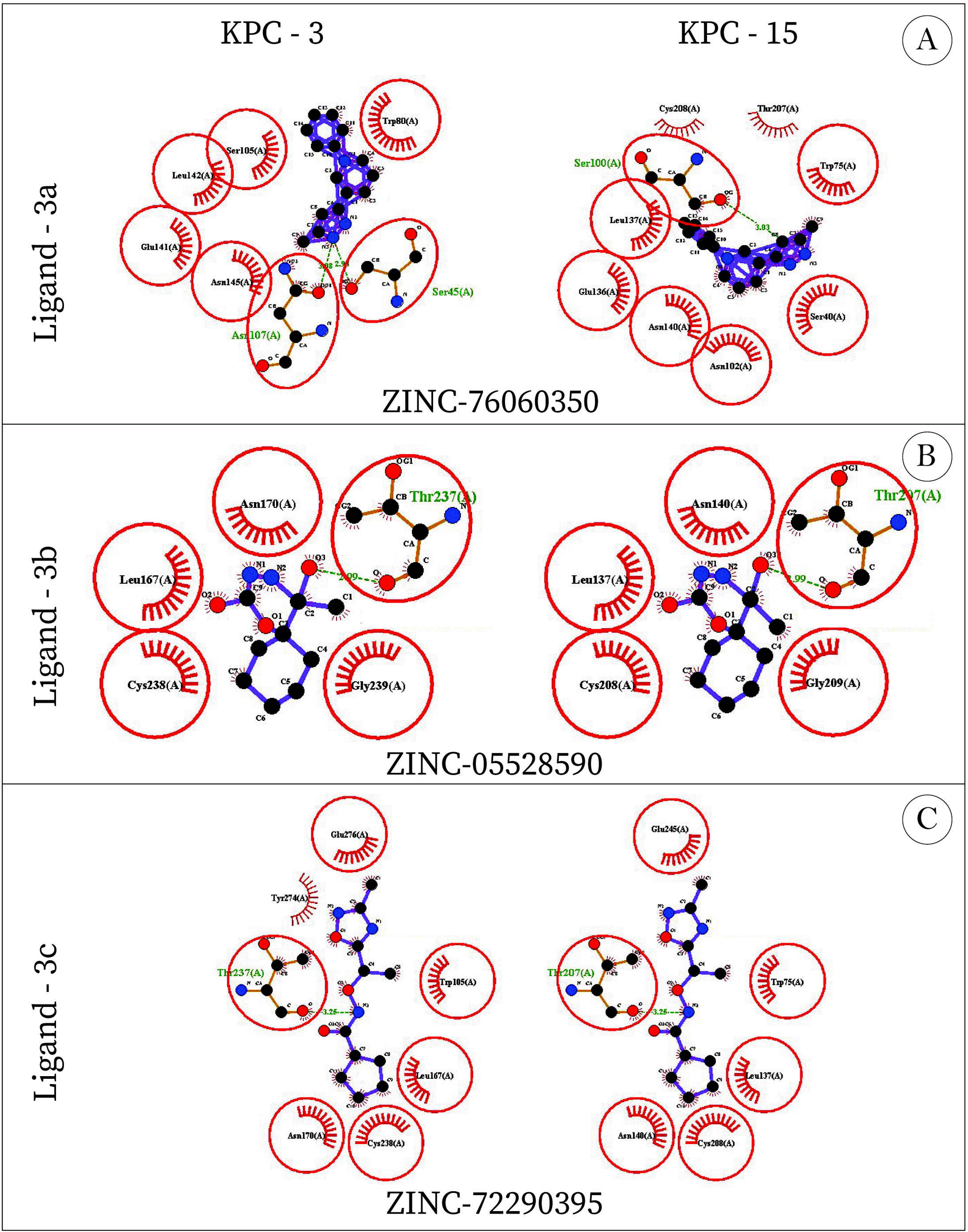
Predicted 2-D structure representation of the interaction of top-3 ligands having combined inhibiting potential of KPC-3 and KPC-15 proteins of K. pneumoniae residue using ligplot+ (A) Ligand-1 (ZINC76060350) docked at the active binding pocket of the KPC-3 protein and KPC-15 protein (B) Ligand-2 (ZINC05528590) docked at the active binding pocket of the KPC-3 protein and KPC-15 protein (C) Ligand-3 (ZINC72290395) docked at the active binding pocket of the KPC-3 protein and KPC-15 protein. Dotted green colour shows H-bonding. The ball-and-stick represents the ligand-protein side chains, where the purple colour shows the ligand bonding with its protein. The non-bonded protein residues interaction with ligand is depicted by spoked arcs. When the two structural models are superposed, the equivalent 3D positions of the potential protein residues are indicated by red circles and ellipses. [37].

**Figure 3:**
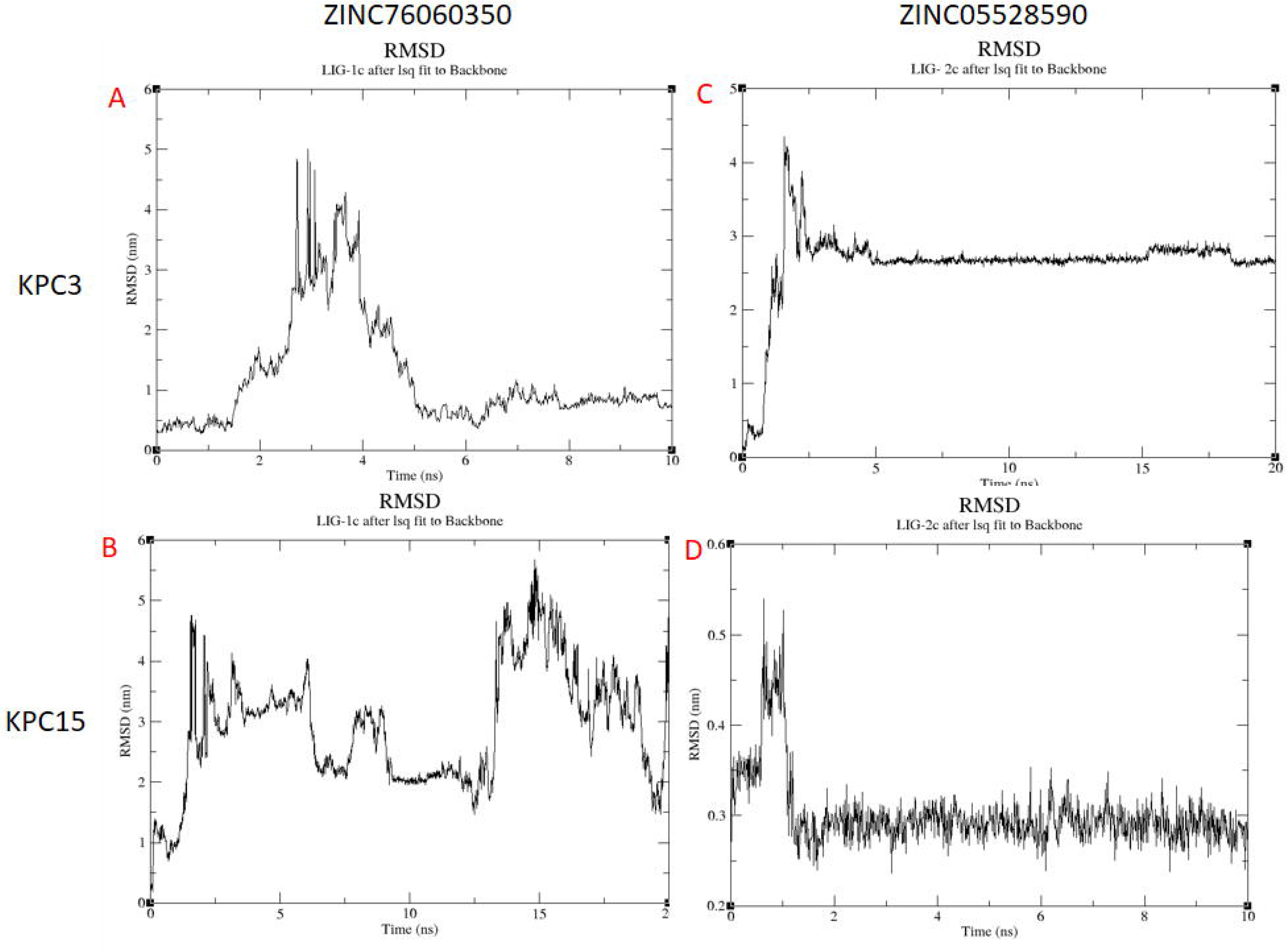
Root-mean-square displacement (RMSD) of the backbone Cα atoms of the KPC3 and KPC15. (A) KPC3 with ZINC76060350 (B) KPC15 with ZINC76060350 (C) KPC3 with ZINC05528590 (D) KPC15 with ZINC05528590

**Figure 4:**
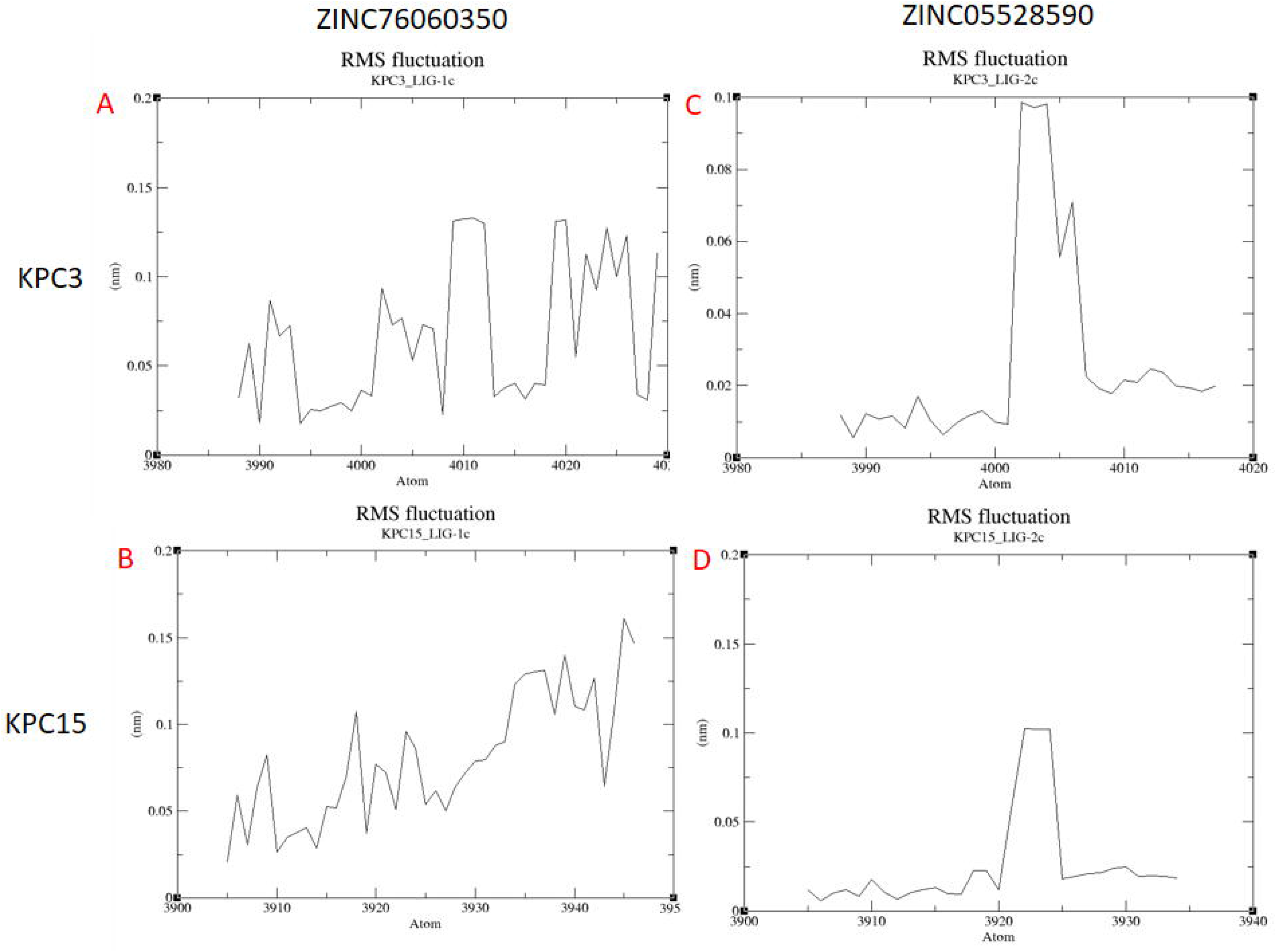
Root-mean-square fluctuation (RMSF) of backbone atoms versus residue number of the KPC3 and KPC15. (A) KPC3 with ZINC76060350 (B) KPC15 with ZINC76060350 (C) KPC3 with ZINC05528590 (D) KPC15 with ZINC05528590

**Figure 5:**
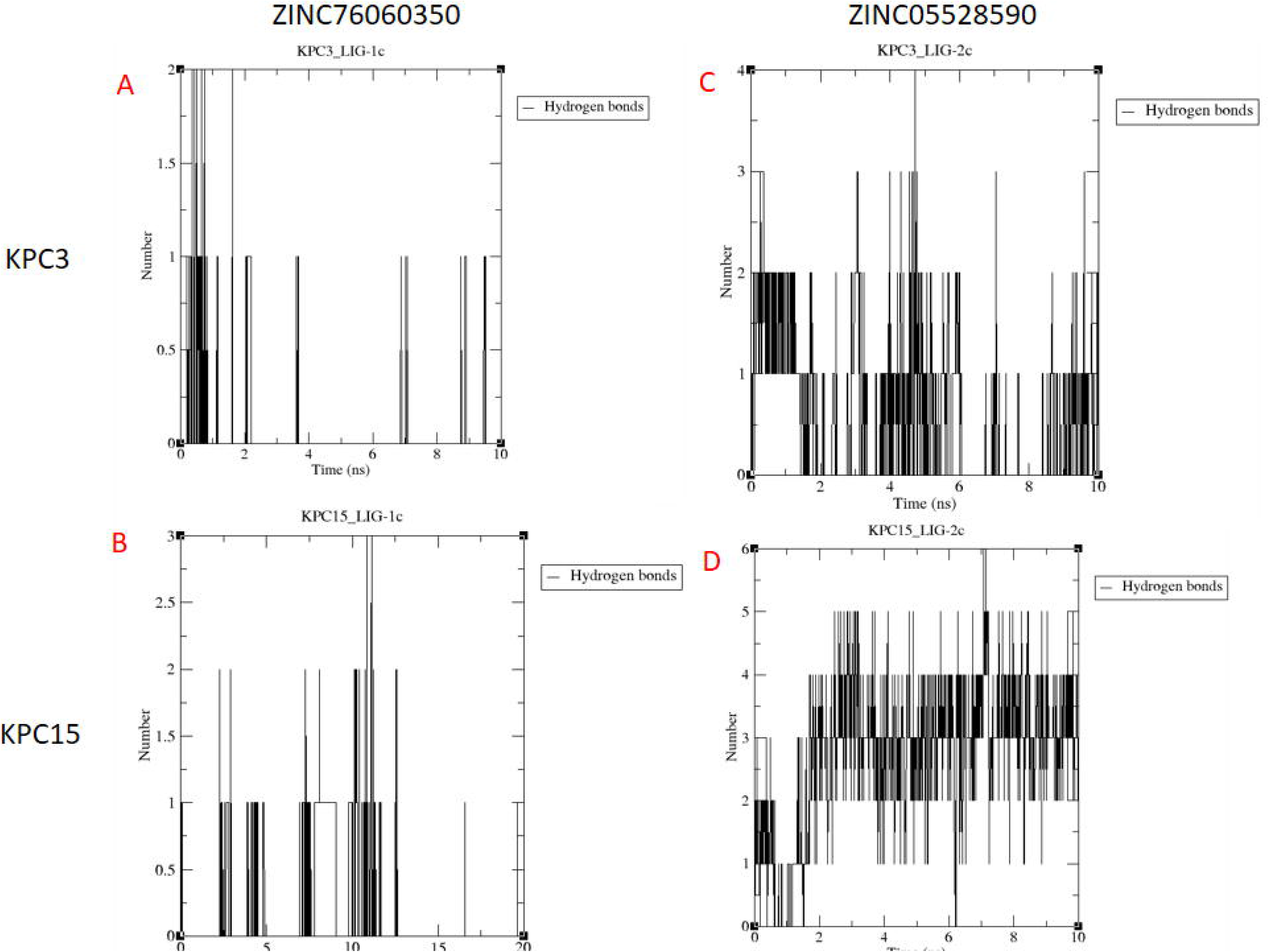
Hydrogen Bond formation of KPC3 and KPC15 residues with ligands (A) KPC3 with ZINC76060350 (B) KPC15 with ZINC76060350 (C) KPC3 with ZINC05528590 (D) KPC15 with ZINC05528590

**Figure 6:**
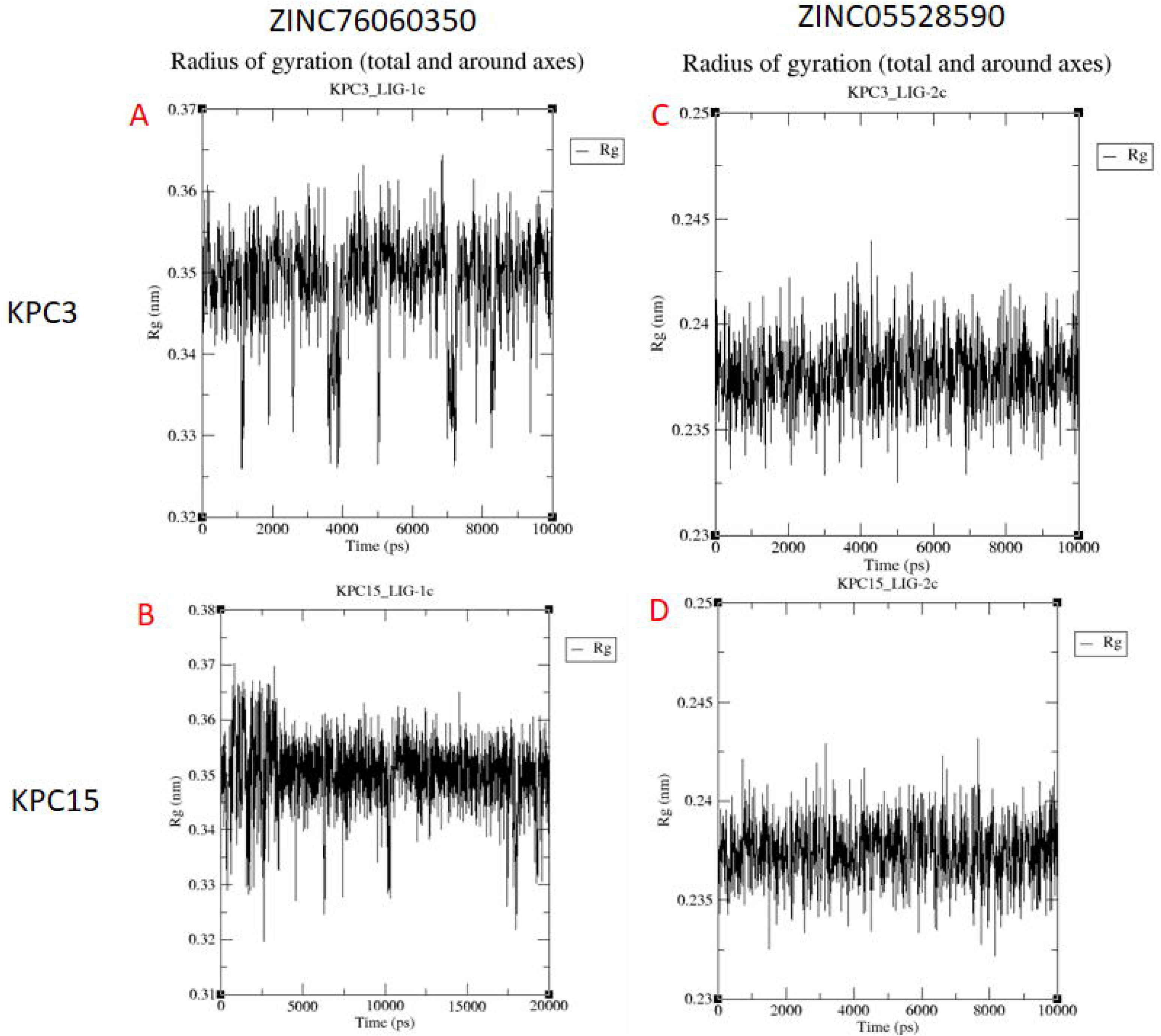
Radius of gyration values (A) KPC3 with ZINC76060350 (B) KPC15 with ZINC76060350 (C) KPC3 with ZINC05528590 (D) KPC15 with ZINC05528590

The analysis of 40 essential ADMET endpoints of all the five potential compounds is listed in table 4. In absorption and distribution, all the top-five compounds can pass through the blood-brain barrier with the maximum permeability probability of 0.99 shown by two compounds-ZINC76060350, ZINC72361341. All five ligands can pass through human intestinal absorption [33] with a minimum probability of 0.92. Good human oral bioavailability was seen with almost all ligands with a maximum probability of 0.89 seen with ZINC71781573. Plasma protein binding shows the probability value of 1.00 by all the ligands, which can be modified by the addition or deletion of the side chain in further stages of the development of the compound. The UGT (UDP-glucuronosyltransferases) were analyzed to investigate the metabolically liable site for lead optimization [34]. Only one ligand showed a UGT catalyzed metabolism- ZINC05528590. There were no interactions seen with estrogen binding receptors and androgen-binding receptors for the majority of the compounds. No hepatotoxicity was seen out of the five compounds. An essential factor to be analyzed during the development of a drug is human ether-a-go-go inhibition, where hERG (the human Ether-go-go-Related Gene) is a gene coordinating the electrical activity of the heart [35]. The compound which has a positive hERG has the potential to cause prolongation of the QT. Two of the five compounds show toxicity against hERG inhibition with a maximum probability value of 0.79. It is not a cause of concern as the computational model of hERG inhibition, and hERG inhibition does not equate to QT prolongation risk, hence for this property confirmation by invitro and invivo studies need to be undertaken [36]. The carcinogenicity potential was not seen in all the five compounds. The biodegradation, eye-corrosion, and eye-irritation are absent for all the compounds.

**Table 4:**
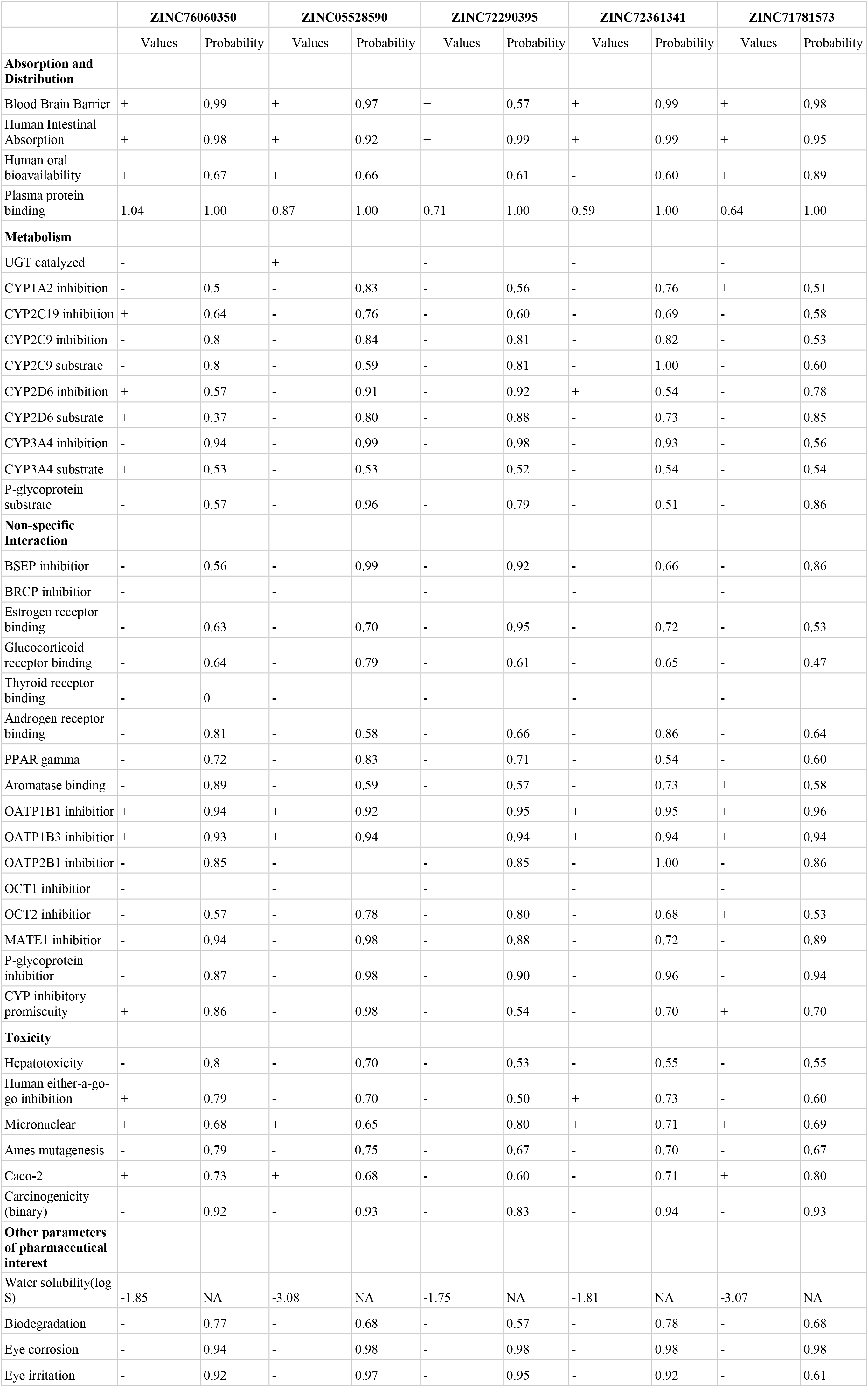
40 ADMET endpoints of the top-five dual inhibitors of KPC-3 and KPC-15, including absorption, distribution, metabolism, non-specific interactions, toxicity and other parameters of pharmaceutical interest, as investigated using admetSAR. NA –stands for Not Applicable

### 3. Structure similarity result

Based on a previous study, Relebactam was identified as a potential inhibitor for KPC-2 showing a dependent relation with imipenem (Papp-Wallace et al., 2018; Zhanel et al., 2018). In-text reference, Relebactam activity against KPC-3 and KPC-15 were not assessed; we checked for the activity of Relebactam against the KPC-3 and KPC-15 and compared the binding energy with that of the binding energy of the top 3 dual inhibitors identified in this study. Our study revealed that ZINC76060350, ZINC05528590, ZINC72290395 gave docking binding energy (Table 3) similar to that of Relebactam (docking energy of Relebactam for KPC-3 and KPC-15 were −6.49 and −6.31, respectively). We then assessed the three identified ligands against Relebactam for structural similarity. (i) The RDKit fingerprint scores between Relebactam and ligand-1c, 2c and 3c were 0.24, 0.22, 0.23 respectively and (ii) Morgan fingerprint score between Relebactam and ligand-1c, 2c, and 3c were 0.26, 0.19, 0.25 respectively. The structure of all three identified ligands is different from Relebactam, showing only 20% similarity (approximately), and therefore, may independently be effective in inhibiting the activity of KPC-3 and KPC-15.

### 4. MD simulation and analysis

RMSD measures the positional change of protein and ligand from the initial simulation stage. The RMSD values of the backbone (protein) in complex with the potential drug candidates were computed concerning the initial complex structure as a reference frame (0 to 10 ns and 0 to 20 ns depending upon the stability attained by the complex). The average RMSD values of ligands ZINC76060350 (ligand 1c) and ZINC05528590 (ligand 2c) in complex with KPC-3 were 0.25, 0.15 and with KPC-15 were 4.65, 0.25 respectively. These values rapidly increase from 1 to 5 ns and 0 to 3 ns, respectively, in ligand ZINC76060350 and ZINC05528590 with KPC-3, while of ligand ZINC05528590 in complex with KPC-15, the value increases from 0 to 1 ns. These rapid increases in peaks and oscillations indicate a binding pocket conformational adaptation of ligand, which later converged to equilibrium state throughout the simulation period. High oscillations were seen throughout the simulation of the ligand ZINC76060350 and KPC-15. In contrast, in the case of ligand ZINC76060350 with KPC-15, it does not attain equilibrium.

The average of RMSF values of the amino acids of KPC-3 and KPC-15 in complex with the ligands were calculated to explore the protein flexibility and residue fluctuations. The average RMSF value for KPC-3 was 0.15 nm throughout the simulation in complex with ligand ZINC05528590, while with ligand ZINC76060350, it was around 0.12 nm. The average RMSF value for KPC15 in complex with ZINC05528590 was 0.12 nm, and with the ligand, ZINC76060350 was 0.1 nm.

Hydrogen Bond formation analysis was performed between the ligands and the protein. Ligand 055 in complex with KPC3 forms more H-bond with binding pocket compared to ligand 760 with KPC-3. In the case of Ligand 055 with KPC-15 forms an average of four H-bond throughout the simulation period

Size and Compactness are the parameters that are associated with the values of the radius of gyration. Initial Rg values of the KPC3-055, KPC3-760, KPC15-055 and KPC15-760 were 0.23 nm, 0.3 nm, 0.23 nm and 1.76 nm respectively. The Rg values of all the protein ligands were at equilibrium throughout the simulation period, proposing that the system has reached the equilibrium state.

#### Limitation and strength of the study

The limitations of the study are that in-vitro or in-vivo validation steps has not been performed and will be a part of different study. The structure of KPC-15 is based on modelled structure rather than the structure determined by spectroscopy. The strength of the study include that we have performed extensive computer aided analysis namely LBVS, SBVS and MD simulation.

## Conclusion

The inhibition of KPC-3 and KPC-15 affects beta-lactamases, which can be used as an advantage for the treatment of infections caused by carbapenemase secreting K. pneumoniae showing resistance to a significant group of antibiotics. According to the results obtained, we suggest that ZINC76060350, ZINC05528590 and ZINC72290395 is the potential dual inhibitors of KPC-3 and KPC-15. The suggested ligands could be taken forward to develop a new drug against a multi-resistant- Klebsiella pneumoniae infection. MD simulations confirmed the mechanism, stability and binding affinity of the ligands with both the protein (KPC3 & KPC15).

